# Differentiation of Alzheimer’s disease based on local and global parameters in personalized Virtual Brain models

**DOI:** 10.1101/277624

**Authors:** J Zimmermann, A Perry, M Breakspear, M Schirner, P Sachdev, W Wen, N.A. Kochan, M. Mapstone, P. Ritter, A.R. McIntosh, A Solodkin

## Abstract

Alzheimer’s disease (AD) is marked by cognitive dysfunction emerging from neuropathological processes impacting brain function. AD affects brain dynamics at the local level, such as changes in the balance of inhibitory and excitatory neuronal populations, as well as long-range changes to the global network. Individual differences in these changes as they relate to behaviour are poorly understood. Here, we use a multi-scale neurophysiological model, “The Virtual Brain (TVB)”, based on empirical multi-modal neuroimaging data, to study how local and global dynamics correlate with individual differences in cognition. In particular, we modeled individual resting-state functional activity of 124 individuals across the behavioral spectrum from healthy aging, to amnesic Mild Cognitive Impairment (MCI), to AD. The model parameters required to accurately simulate empirical functional brain imaging data correlated significantly with cognition, and exceeded the predictive capacity of empirical connectomes.

## 1. Introduction

The cognitive and anatomical changes that occur in dementia due to Alzheimer’s disease (AD) have been widely documented (Braak and Braak, 1991; Braak et al., 1993; Hyman et al., 2012; Van Hoesen and Damasio, 1987; Weiner et al., 2012). Changes in structure and function occur across a range of spatial scales. Neurofibrillary tangles (NFT) disrupt axonal flow via tau phosphorylation, which in turn disrupt functional communication. The accumulation of tau protein results in the degeneration of axonal tracts, causing further damage in global functional connectedness, and finally, neuronal death.

More recently, neuronal hyperactivity has been implicated in the degenerative cascade of AD. This hyperactivity, characterized by changes in local excitatory and inhibitory neuronal populations as well as global hyperexcitation, have been documented in both Mild Cognitive Impairment (MCI) and AD (Busche and Konnerth, 2015; Celone et al., 2006; Dickerson et al., 2005; Jones et al., 2016; Sperling et al., 2010). The imbalance in excitatory and inhibitory neural circuits disrupts hippocampal functioning, which likely leads to cognitive decline (Goutagny and Krantic, 2013; Scott et al., 2012; Verret et al., 2012). The change in excitation/inhibition is particularly intriguing, as it may be causally linked to the amyloid buildup that is a regular characteristic of AD (Gleichmann et al., 2011; Gleichmann and Mattson, 2010; Palop and Mucke, 2010). Interestingly, the excitation that characterizes some subtypes of AD has actually been linked to an increased risk of seizure (Amatniek et al., 2006; Mark et al., 1995). Local excitation, integrated across the network via inter-area connections increases the coordinated activity of the network via heightened FC (Deco et al., 2014b), which is a feature of seizure activity (Mishra et al., 2013).

At the global level, AD has been described as a disconnection syndrome, characterized by the degeneration of the connectome and of network organization that is governed by these long-range connections (Delbeuck et al., 2003; He et al., 2008; Sanz-Arigita et al., 2010; Stam et al., 2007; Supekar et al., 2008; Wang et al., 2007). Both structural and functional connectivity (SC, FC) are affected (Balachandar et al., 2015; Filippi and Agosta, 2011; Matthews et al., 2013; Soldner et al., 2012; Sun et al., 2014). Changes in global network organization have been reported as a result (Delbeuck et al., 2003; He et al., 2008; Sanz-Arigita et al., 2010; Stam et al., 2007; Supekar et al., 2008; Wang et al., 2007).

To date, linking findings of AD pathology at these different scales of interrogation has proven elusive. Computational models are a promising strategy for combining network-level connectivity with neural mass models. These models allow for the systematic exploration of the optimal levels of neural parameters such as excitation, inhibition, coupling or conduction delays between distant regions. Computational models have already proved informative for understanding the link between SC and FC in the brain, for example via lesion studies (Alstott et al., 2009; Honey and Sporns, 2008). Several studies have focused on modeling changes in interconnected excitatory and inhibitory neural populations as they relate to network organization in clinical populations (de Haan et al., 2012; de Haan et al., 2017; Yang et al., 2016). Two of these studies have documented such changes in Alzheimer’s disease. In the first study, De Haan et al. (2012) emphasized the pathological effects of hyperexcitability by showing that the vulnerability of hub regions in AD is related to their increased excitability. De Haan et al (2017) showed that, contrary to their hypothesis, selective stimulation of excitatory neurons resulted in the prolonged preservation of oscillations, connectivity and network topology. The results of the study emphasize the unpredictability of excitation/inhibition on the complex dynamics of the brain. However, neither of these studies linked these changes to observed differences in cognition, as they were more interested in the influence on network properties.

The goal of the present work was to elucidate whether optimal functioning points of local (excitation, inhibition) and global (long-range coupling, conduction velocity) brain dynamics can be linked to individual differences in cognition from healthy aging, to MCI, to AD. Our objective was to understand whether variability in these parameters can be informative for individual differences in AD. To this end, we used the novel neuroinformatics platform “The Virtual Brain” (TVB, thevirtualbrain.org), a large-scale simulator of brain dynamics based on connectivity metrics with the ability to target individual neural populations (Jirsa et al., 2017; Jirsa et al., 2010; Ritter et al., 2013; Roy et al., 2014; Sanz Leon et al., 2013; Sanz-Leon et al., 2015; Woodman et al., 2014). Here, we modeled changes in interconnected excitatory and inhibitory neural populations that occur at the local level along with global connectivity changes. Each brain region was modelled as a local population comprising connected excitatory and inhibitory neurons linked with NMDA and GABA synapses (Deco et al., 2014a). Neural activity was generated as a function of the intra-areal local parameters – the excitatory and inhibitory inputs, and the inter-regional long-distance individual diffusion-derived SC that integrates these across the network (Deco et al., 2014b).

The strength of TVB is that it is based on *individual* structural and functional data and thus the parameter values obtained reflect personalized Virtual Brains (Ritter et al., 2013; Sanz-Leon et al., 2015; Woodman et al., 2014). We can compare these parameters across individuals and study their association with individual differences in cognition. Previous work has shown that biophysical parameters derived from TVB modeling can describe healthy neural dynamics (Jirsa et al., 2010; Kunze et al., 2016; Roy et al., 2014), as well as disease. As a proof of concept, it has been shown that these biophysical parameters correlate with motor recovery from stroke (Adhikari et al., 2015; Falcon et al., 2016a; Falcon et al., 2016b; Falcon et al., 2015), and the generation of epileptic seizures (Jirsa et al., 2014), along with seizure progression (Jirsa et al., 2017). The biophysical parameters derived from TVB modeling have been shown to correlate with a variety of clinical phenotypes, and as such offer a potential for translation to clinical applications.

In the present study, we sought to identify biophysical model neural parameters that associated with cognition along the spectrum from healthy controls to MCI to AD. We include non-demented amnestic MCI (aMCI) subtypes within our investigation, given these individuals with memory-based impairments demonstrate increased rates of progression to AD – and are assumed to be experiencing a transitory, pre-clinical stage (Fischer et al., 2007; Petersen et al., 2014; Ward et al., 2013). aMCIs show widely variable rates of conversion to AD (Ward et al., 2013), and thus differences within this group are of interest to the present study. Importantly, we also create several models that reflect the progression of AD. Neurodegeneration and functional changes occur first in limbic and temporal regions of the brain, and later in motor and sensory areas (Braak and Braak, 1991; Braak et al., 1993). The pattern of destruction follows the clinical phenotype; memory is targeted first, followed only later by sensory and motor function. Accordingly, modeling is performed on a limbic subnetwork (Limbic SubNet model), as well as the full brain (Whole Network model). Any discrepancies between the Limbic SubNet and the Whole Network model may be characterized along the spectrum of cognitive phenotypes from healthy, to aMCI to AD.

## 2. Methods

### 2.1 Subjects

Comprehensive behavioral data were acquired for 124 participants from the fourth wave of the Sydney Memory and Ageing study (MAS) (Perminder S. Sachdev et al., 2010; Tsang et al., 2013). At study baseline (approximately six years prior to acquisition of the current data), these community-dwelling participants were initially between 70-90 years of age. Subjects were stratified based on cognitive performance into three groups: 73 healthy controls (HC), 35 with amnestic MCI (aMCI), and 16 Alzheimer’s Disease (AD).

For all study participants, exclusion criteria at baseline included a Mini-Mental Statement Examination (Folstein et al., 1975) adjusted score below 24, diagnosis of dementia, developmental disability, neurological or psychiatric diseases, or inadequate comprehension of English to complete a basic assessment. The study was approved by the Human Research Ethics Committee of the University of New South Wales, and participants gave written, informed consent.

### 2.2 Neuropsychological measures

Twelve individual neuropsychological tests were administered to cover a broad range of cognitive functions. These domains assessed included attention/processing speed, memory, language, visuospatial ability, and executive function. These tests were grouped into cognitive domains as part of the boarder longitudinal study (MAS) (Kochan et al., 2010; P.S. Sachdev et al., 2010) – and selected accordingly to the primary cognitive function they assess (See Table S1). This was based upon the extant literature and the widespread practice used by neuropsychologists (Lezak, Howieson, & Loring, 2004; Strauss, Sherman, & Spreen, 2006; Weintraub et al., 2009). Further rationale for the cognitive groupings and their scale-items homogeneity estimates are outlined elsewhere (A. Perry et al., 2017).

Other measures such as the MMSE and the National Adult Reading Test (NART IQ) (Nelson & Willison, 1991) were also administered. The NART, which was administered to a subset of the current population at study baseline, estimates premorbid intelligence levels (Bright, Jaldow, & Kopelman, 2002).

The neuropsychological assessments were used in two different ways:

#### a Diagnostic purposes

The neuropsychological and clinical profiles of the study participants determined the individual’s classification into one of the three population groups (Perminder S. Sachdev et al., 2010; Tsang et al., 2013).

*Healthy Controls* (HC): Performance on all neuropsychological test measures higher than a threshold of 1.5 SDs below normative values (matched for age and education level). The criteria for selection, and the demographic matching that was used to establish a normative reference is described in full elsewhere (Tsang et al., 2013).

*Mild Cognitive Impairment* (MCI): If the following international consensus criteria (Winblad et al., 2004) of MCI were met:

1. Subjective complaint of memory decline or other cognitive function (from either the participant or informant).
2. Evidence of cognitive-decline, derived here by performance 1.5 SDs below the normative values on any of the neuropsychological tests.
3. Normal or minimally impaired functional activities, determined by informant ratings on the Bayer-ADL scale (Hindmarch et al., 1998).
4. No current diagnosis of Dementia according to DSV-IV criteria (APA, 2000).

Only MCI individuals demonstrating memory-based impairments - hence meeting the criteria for the amnestic MCI (aMCI) (Petersen, 2004) subtype - were included in the current study. Fifteen of these aMCI individuals were further representative of the multi-domain aMCI subtype (md-aMCI) (Petersen, 2004), whom are characterized by additional impairments in non-memory domains.

*Alzheimer’s Disease* (AD) *patients*: A diagnosis of Alzheimer’s Disease according to DSM-IV criteria (APA, 2000) – according to a clinical expert panel comprising of geriatric psychiatrists, neuropsychiatrists, clinical neuropsychologists and clinical psychologists. All clinical and structural MRI data (where available) were used in the diagnostic decision.

#### b Correlative metrics with model parameters

Performance on the individual test scores were transformed into quasi Z-scores, based upon the mean and SDs for a healthy reference group (N = 723), identified from all the study participants at baseline (approximately 6 years prior to acquisition of the current wave’s data). Domain scores were calculated as the average of the transformed test scores comprising each domain. The only exception was the visuospatial domain that was represented by a single measure. The memory domain composite was further subdivided into verbal memory after exclusion of a visual retention test (Benton, Sivan & Spreen, 1996). If necessary, the signs of the z-scores were reversed so that higher scores reflect better performance. The correspondence between neuropsychological cognitive domain z-scores and clinical diagnosis classification (healthy control, MCI, AD) is reported in the Results.

### 2.3 Imaging data

Structural, diffusion (dMRI) and resting-functional MRI (rs-fMRI) data were acquired on a Phillips 3T Achieva Quasar Dual scanner. dMRI were acquired with a single-shot echo-planar imaging (EPI) sequence with the following parameters: TR = 13,586 ms, TE = 79 ms, 61 gradient directions (b = 2400 s/mm^2^), a non-diffusion weighted acquisition (b = 0 s/mm^2^), 96×96 matrix, FOV = 240 x 240 mm^2^, slice thickness = 2.5mm, yielding 2.5mm isotropic voxels. For rs-fMRI, participants were instructed to lie quietly in the scanner and close their eyes. A T2* weighted EPI sequence with the following parameters was acquired: acquisition time = 7:02, TE = 30 ms, TR = 2000 ms, flip angle = 90º, FOV 250 mm, 136 x 136 mm matrix size in Fourier space. 208 volumes were acquired, each consisting of twenty-nine 4.5 mm axial slices. A structural T1-weighted image was also acquired with the following parameters: TR = 6.39 ms, TE = 2.9 ms, flip angle = 8°, matrix size = 256 × 256, FOV = 256 × 256 × 190, slice thickness = 1 x 1 × 1 mm isotropic voxels.

Data from all imaging modalities was then inspected in FSL View (Smith et al., 2004) for quality checking purposes. Subjects were removed if any of their scans had artifact issues, including slice dropouts on the diffusion-images (defined by zebra-like blurring or complete dropout) (Pannek et al., 2012b), complete orbitofrontal EPI signal dropout (Weiskopf et al., 2007), ringing on T1-images, or severe geometric warping.

### 2.4 Preprocessing and tractography of diffusion data

Steps involving the preprocessing and whole-brain tractography of dMRI data were similar to those performed for a subset of the current healthy control population (Perry et al., 2015b; Roberts et al., 2017). In short, head motion correction was performed by rotation of the gradient directions (Leemans and Jones, 2009; Raffelt et al., 2012), and spatial intensity inhomogeneities were reduced via bias field correction (Sled et al., 1998).

Estimates of fibre orientation and subsequent whole-brain tractography were performed within MRtrix3 (v0.3.12-515; https://github.com/MRtrix3) (Tournier, Calamante & Connelly, 2012), Fiber orientation distribution (FOD) functions were first estimated using constrained spherical deconvolution (CSD) (lmax = 8) (Tournier et al., 2008) of the diffusion signal in voxel-populations with coherently-organized (FA > 0.7) fiber bundles. We then performed iFOD2 (Tournier, Calamante & Connelly, 2010) probabilistic tracking to propagate 5 million fiber tracks from random seeds throughout the brain for each subject, with the following parameters: step size = 1.25 mm, minimum track length = 12.5 mm, maximum length = 250 mm, FOD threshold = 0.1, curvature = 1 mm radius.

The sequences were acquired when reverse phase-encoding direction approaches were not the standard procedure within acquisition protocols. Given that the alignment between the diffusion and anatomical priors will not be perfectly accurate, seeding of the tractograms was not performed upon the grey/white-matter interface (i.e. ACT) (Smith et al., 2012), nor were streamline-filtering approaches applied (i.e. SIFT/SIFT2) (Smith et al., 2013).

### 2.5 fMRI processing

fMRI data processing was performed using the Data Processing Assistant for Resting-State fMRI (DPARSF, v 3.2) (Chao-Gan and Yu-Feng, 2010), which calls functions from SPM8 (http://www.fil.ion.ucl.ac.uk/spm/). Slice-timing correction (realignment to mean functional image) was performed, followed by co-registration to the structural image (via 6 DOF). We then conducted linear detrending, nuisance regression of 24 motion parameters (Friston et al., 1996) and segmented WM/CSF signals (Ashburner & Friston, 2005). An average structural brain template across all participants was generated (DARTEL) (Ashburner, 2007), and native functional images were then transferred to MNI space (3 mm) via this template. Lastly, smoothing (8 mm) and temporal band-pass filtering (0.01-0.08 Hz) was performed. Full description of the steps involved for the pre-processing of these data are provided elsewhere (Perry et al., 2017)

### 2.6 Functional and structural brain networks

Connectomes representing patterns of SC and FC were constructed for all subjects, estimated from streamline-tractography maps and BOLD rs-fMRI signals, respectively. The widely-used AAL parcellation (within MNI space) was used here to derive the inter-areal connectivity estimates (Tzourio-Mazoyer et al., 2002). The choice of parcellation was motivated by previous work. The performance of identifying atrophy patterns in Alzheimer’s using the standard AAL template has been compared with a subdivision of the AAL parcels into regions (N = 487) of smaller size (Mesrob et al., 2008). The AAL template indeed demonstrated higher classification accuracy when using inter-cohort validation, relative to the finer-grained parcellation (Mesrob et al., 2008). The AAL template has also been widely used in AD studies (Savio, 2017; Tijms et al., 2018; Wang et al., 2006), and was recently used within a whole-brain computational modeling approach to elucidate AD-related characteristics such as amyloid beta and tau (Demirtas et al., 2017). Lastly, the AAL template was further chosen in lieu of the FreeSurfer parcellation because the noted acquisition procedure circumvents the imperfect alignment between the diffusion and T1-weighted imaging modalities.

For the structural connectomes, parcellations in subject-space were first achieved with FSL5 (Smith et al., 2004) by linearly co-registering each individual FA image into the FMRIB standard space template. By applying the inverse of this above transformation matrix, the AAL parcellation was subsequently transformed into subject-space. Given that the functional images were already within MNI space, the parcellation template was not transformed to native-space for functional network construction.

FC was computed as the Pearson’s correlation coefficient of the mean BOLD signals (i.e. all region *i* voxels) between all *ij* region pairs. For the SC matrices, two matrices were computed: (1) a weights connection matrix, and (2) a distance matrix used to assess conduction delays in the model. In the weights matrix, *Wij* represented the total number of streamlines that start/terminate within a 2mm radius of regions *i* and *j*. Note that the diffusion signal becomes noisier and weaker around the grey-white matter boundaries. By selecting a 2mm radius (default parameter within MRtrix tck2connectome), we ensured these erroneous fiber terminations were still identified as connectome edges. Whilst this streamline identification approach could potentially lead to over-sampling of the streamline weights, we otherwise circumvent the potential for false negatives and under-sampling within the connectome edges. *W*_*ij*_ were corrected by the euclidean distance between *i* and *j* and also total volumetric size (summed number of voxels) of the two regions. A subset of the AAL regions was used in the Limbic SubNet model, which characterized our AD subnetwork (See Supplementary Table S2 for a region list).

### 2.7 Quality control of SC

We implemented stringent diffusion quality control procedures, which have been standard protocol for our other published works (Perry et al., 2015a; Roberts et al., 2016). In particular, each raw (i.e. unprocessed) diffusion volume of every subject was visualized. Any participant with notable motion effects and/or severe spatial distortions was removed from the study, operationalized by either complete signal dropout and “zebra-like” blurring of slices (Andersson and Sotiropoulos, 2016; Pannek et al., 2012a). Furthermore, each subject’s Fractional Anisotropy image was visualized, as was the accuracy of the co-registration between the FA image and AAL parcellation (that was originally within FMRIB FA space). The FMRIB FA template was chosen for an improved registration with each subject’s FA image. The FMRIB FA template was based upon normalization of independent FA images, and relative to the MNI T1 brain provided with FSL, provided a more similar anatomical shape to the subject FA images. An example of the alignment accuracy between the FMRIB and subject’s FA image are provided in Figure S1D.

The steps involved in the estimates of fibre trajectories and the subsequent construction of each subject’s SC are summarized within Figure S1. Note that SC weights were exponentially distributed (See Figure S2), and that SC networks were not disconnected as we have used a dense-seeding approach with the tractography, and a relatively coarse parcellation scheme. Moreover, thresholding was not performed. To identify this did not potentially influence our results, we 1) checked that raw connectome density distributions were consistent across clinical groups (ANOVA: *F*(2,121) = 0.41, *p* = 0.66) and 2) that density did not correlate with cognitive performance (*r* = −0.039, *p* = 0.67). The parcellation and affine co-registration (to the diffusion images) algorithm were identified to yield an acceptable standard of alignment accuracy across the population groups (Figure S1F).

### 2.8 Computational modeling with TVB

The Virtual Brain is a multi-scale modeling approach that combines local parameters (i.e. population excitation, inhibition) with global long-range parameters that take into account the connectivity structure (i.e. global coupling) and dynamical interactions between regions (i.e. conduction velocity). The TVB modeling process is as follows: 1) incorporation of subject SC matrices (weights and track distances); 2) selection of a local model and parameters (See Section 2.8.1 below for details); 3) selection of global parameters (i.e. conduction velocity, global coupling); 4) simulation of rsfMRI time series based on an integration of global and local dynamics; 5) computation of the simulated rsFC and fitting to the empirical rsFC; 6) iterative optimization by re-running steps 1-5 until an optimal fit is achieved; and 7) correlating optimal model parameters with patient phenotype (i.e. cognitive scores). See Figure 1 for a summary. This process was described in detail elsewhere (Ritter et al., 2013; Sanz-Leon et al., 2015; Woodman et al., 2014).

**Figure 1.**
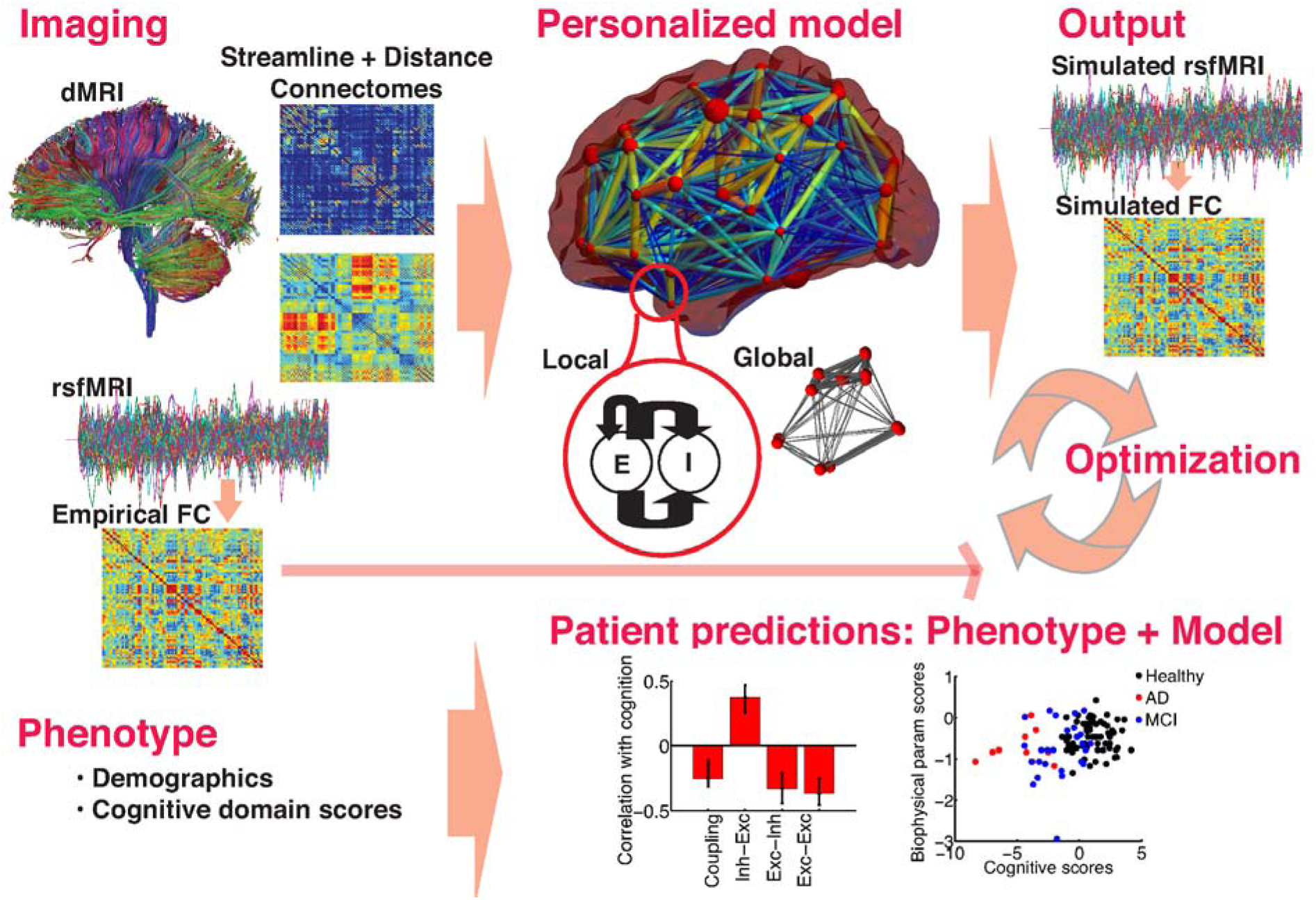
Workflow in TVB Modeling: Graphical representation of the sequential steps taken in this study: a) Individual empirical neuroimaging data; dMRI is the input to the model, rsfMRI is used for model fitting and optimization, b) Personalized model: integration of global and local dynamics; c) Output: simulated rsfMRI time series, FC is then computed, d) Optimization: model is optimized by iterative adjustment of parameters to achieve best fit of the simulated to the empirical FC. e) Phenotype: individual patient demographics, cognitive domain scores etc, these are used to make, f) Patient predictions, by correlating cognitive domain scores with model parameters.

#### 2.8.1 Model

We used the reduced Wong Wang model (Deco et al., 2014b), a mean field model that simulates local regional activity via interconnected populations of excitatory pyramidal and inhibitory neurons, with NMDA (excitatory) and GABA-A (inhibitory) synaptic receptors (Deco and Jirsa, 2012; Deco et al., 2014b; Wong and Wang, 2006). The model is based on firing rates and synaptic gating activity. The model was chosen as it is one of the more refined local models with an ability to model the balance between excitation and inhibition, which is disrupted in AD (Busche and Konnerth, 2015; de Haan et al., 2012; de Haan et al., 2017; Palop et al., 2007; Verret et al., 2012). As with all populations models of large-scale neuronal activity, the model rests upon a combination of abstraction and mean-field dimension reduction (Breakspear, 2017). This is required to render it tractable, to improve parameter identifiability and to speed up computation time, hence facilitating parameter exploration and optimization. The Wong Wang model retains local within compartment excitatory and inhibitory connectivity, mediated by local synaptic currents (Wong and Wang, 2006). These form part of our central hypothesis. The model is of comparable complexity to other models used in computational assays of healthy and compromised brain dynamics (de Haan et al., 2017), being more complex than reduced oscillator models that have also been employed (Breakspear et al., 2010), whilst slightly less complex than population models that retain multi-layer connectivity (Bastos et al., 2012). Note however, that we do not test layer-specific hypotheses. All population models are by definition more abstract than multi-compartment spiking neural models: while these play an important role in dissecting microscopic models of disease, their high dimensionality and myriad of parameters challenges attempts to scale then to understand network and whole brain dynamics (Breakspear, 2017).

Each local population forms a node, which are then coupled together through subject-specific connectomes to model the corresponding whole brain network dynamics. The behaviour at each node, *i*, is defined according to a set of stochastic differential equations (Deco et al., 2014b) describing synaptic currents *I*, synaptic gating variables, *S* and firing rates *r*. Currents in excitatory 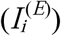 and inhibitory populations 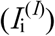 at node *i* are modelled as a combination of local recurrent feedback arising from the local fraction of activated synapses (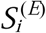 and 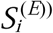), plus input from excitatory neurons in distant nodes 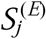. These are given as,

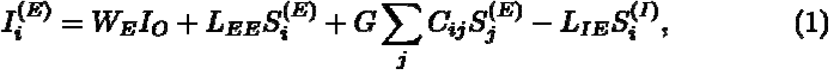

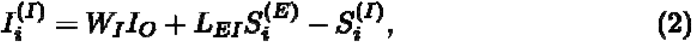

where *W*_*E*_ and *W*_*I*_ scale the effect of a fixed external current *I*_*O*_ (= 0.382). L_EE_ scales the local excitatory recurrence (excitatory influences on excitatory populations, E-E) and *L*_*IE*_ scales the local inhibitory synaptic coupling (inhibitory influences on excitatory populations). *L*_*EI*_ scales the local excitatory synaptic coupling (excitatory influences on inhibitory populations, E-I). External input from other nodes is conveyed through the subject-specific connectome *C*. *G* is the global coupling scaling factor for *C*_*ij*_, the structural connection matrix weights of tracks between regions *i* and *j*. Note that *L*_*EE*_, *L*_*IE*_, and *L*_*EI*_ correspond to the combined influence of *w+* and *Ji* or *JNMDA* that are described elsewhere (Deco et al., 2014b). The firing rates of the neural populations in node *I* are then described by passing the corresponding input currents through a sigmoid-shaped activation function *H*,

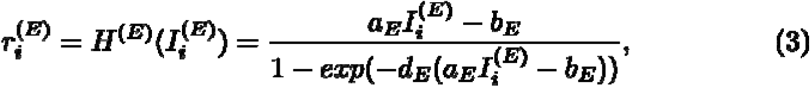

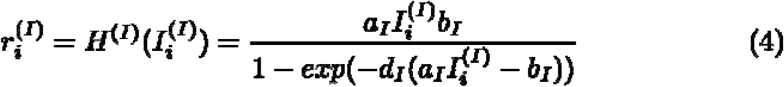

where 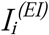 is the input current to the (excitatory or inhibitory) population *i*. The synaptic gating constants are *a*_*E*_ = 310 (nC^-1^), *a*_*I*_ = 615 (nC^-1^), *b*_*E*_ = 125 (Hz), *b*_*I*_ = 177 (Hz), *d*_*E*_ = 0.16 (s), *d*_*I*_ = 0.087 (s). The set of equations are then closed by expressing the temporal evolution of the simulated excitatory and inhibitory synaptic gating variables *S* as a function of the corresponding firing rates,

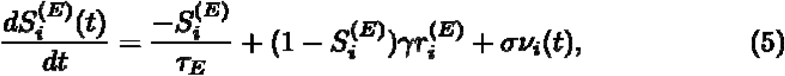

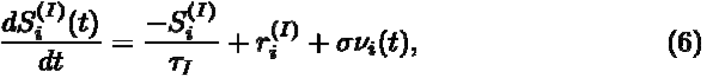

Note that in the absence of input, these variables relax back to zero with time constants for the decay of synaptic gating variables of NMDA and GABA receptors given by *τ*_*E*_ = *τ*_*NMDA*_ = 100 (ms) and *τ*_*I*_ = *τ*_*GABA*_ = 10 (ms). The constant *γ* = 0.000641 scales the self-feedback of local firing. Stochastic influences are introduced to these self-consistent equations via *ν*_*i*_, an uncorrelated Gaussian noise source with unit standard deviation scaled in amplitude by *σ* = 0.01 (nA). The model parameter values are provided in Table S3.

#### 2.8.2 Model fitting

Parameter space exploration was performed so that local and global parameters were adjusted iteratively to optimize model fit to the empirical data. Parameter fitting is done across the whole network, so that parameters are consistent across all nodes in the modeled network.

##### Global parameters

Model fits were maximized by varying conduction velocity and global coupling. Conduction velocity reflects the speed of signal transfer across white-matter fibers (in m/s). It is derived from the Euclidean distances. Previous studies have suggested the importance of conduction delays for resting-state BOLD (Deco et al., 2009; Ghosh et al., 2008). Global coupling (*G*, arbitrary units) is a scaling factor for the anatomical weights. It determines the balance between global long-range input from distal regions, and the local recurrent input from the local neural populations within regions. High global coupling denotes greater weighting of the global over the local input, such that activity in node *i* is driven more by activity propagated over the long-range connections from other nodes *j*.

##### Local parameters

The most salient group of parameters at the local level determine the coupling between excitatory and inhibitory neuronal populations. This is modeled via 3 local parameters: excitation from recurrent excitatory-excitatory populations (*L*_*EE*_), excitatory input to inhibitory populations (*L*_*EI*_), and dampening of activity via recurrent inhibition from inhibitory to excitatory populations (*L*_*IE*_).

A progressive parameter space exploration was performed, with parameters varied one at a time in the following order: conduction speed, global coupling (*G*), inhibition-excitation (*L*_*EI*_), excitation-inhibition (*L*_*EI*_), excitation-excitation (*L*_*EE*_), in a manner similar to Falcon et al., 2015, 2016. The following parameter ranges were used ([min, max, step size]: global coupling ([0.5, 2.0, 0.1]), inhibition-excitation ([0.4, 2.6, 0.1]), excitation-inhibition ([0.025, 0.5, 0.025]), excitation-excitation ([0.5, 2.0, 0.1]). Note that parameter ranges were chosen based on parameters ranges in Deco et al., 2014.

Parameter exploration was performed according to the following heuristic: For each set of parameter combinations, individual resting-state BOLD fMRI time series of duration 3 minutes were simulated using a Balloon-Windkessel hemodynamic model (Buxton et al., 1998; Friston et al., 2000). Resting-state BOLD FC was then derived from each of the simulated time series. Specifically, model fitting was achieved by iterative optimization of the biophysical parameters until the maximum correlation to our subject-specific empirical FC across the range of input parameters was obtained. Pearson’s correlations between the lower triangle of the empirical and the simulated FC matrices were used as the fitting criterion. Optimal biophysical parameters for each subject were those that achieved the best fit to the empirical FC.

### 2.8.3 Whole network, Limbic SubNet, Embeddedness of Limbic SubNet

As not all brain regions are affected simultaneously in AD, model parameters were estimated via three different approaches:

1. Whole Network modeling: Model input is the full network SC derived from the AAL parcellation. Simulated BOLD FC was derived for all regions.
2. Limbic SubNetwork: As certain limbic and temporal regions show the greatest neuropathological changes and degeneration early in the disease (Braak and Braak, 1991; Braak et al., 1993; Van Hoesen and Solodkin, 1994), associated with memory declines (Braak and Braak, 1991; Braak et al., 1993), we did additional modeling in a subnetwork that included the following regions in both hemispheres: cingulum (anterior, middle, posterior), hippocampus proper, parahippocampus, amygdala, temporal pole (superior, middle) (See Table S2). Model input was an SC of this subnetwork, derived as a subset of the AAL parcellation. Simulated BOLD FC of these regions was derived.
3. Embeddedness of Limbic SubNet: In order to characterize the discrepancies of brain dynamics in the Limbic SubNet and the larger network, we quantified the discrepancy between Limbic SubNet and Whole Network model parameters (Limbic SubNet – Whole Network optimal parameters).

### 2.9 Statistical analysis

Analyses were done by comparing individuals’ optimal model parameters: (1) with neuropsychological cognitive z-scores in a continuous scale and (2) between groups (healthy, aMCI, AD - based on clinical diagnosis). See Figure S3 for correspondence between cognitive scores and clinical diagnosis groups.

#### (1) Model correlates of cognition

To investigate the relationship between individual z-scores on the 6 cognitive domain measures (attention, language, executive functioning, visuospatial, memory, verbal memory), and the optimal biophysical parameters across subjects, we conducted a set of Partial Least Squares (PLS) correlation analyses (Krishnan et al., 2011; McIntosh and Lobaugh, 2004) Behavioural PLS is comparable to a Canonical Correlation Analysis, such that it decomposes the correlation between two variables into latent variables (LVs), or components, which identify the maximum least squares relationship between the variables. PLS is more robust than canonical correlation when there is potential for colinearity amongst the measures which is often the case in neuroimaging data. The LVs in PLS are comparable to LVs in canonical correlation.

The 4 biophysical model parameters were inputted as X and the 6 cognitive performance domains were inputted as Y, and the correlation between the two was computed. The cross-block correlation matrix was decomposed with SVD, yielding mutual orthogonal LVs, which have weights for X and Y variables and the singular value (i.e., covariance) that conveys the magnitude of the relation between X and Y for that LV. Within each LV, the reliability of the contributions of each measure in X and Y were captured via the bootstrap estimation of confidence intervals for the weights converted to correlations. Permutations tests were used to assess the statistical significance of each LV, where its covariance (singular value) is compared to a distribution of random permutation of data pairings (McIntosh & Lobaugh, 2004).

#### (2) Group differences in biophysical model parameters

Differences in model parameters across clinical groups (healthy controls, MCI, AD) and network model (Limbic SubNet, Whole Network) were assessed using a non-rotated task PLS with two factors. Non-rotated task PLS is comparable to a MANOVA in that it tests for differences between groups with several dependent variables. However, non-rotated PLS is more robust than a MANOVA when there is a potential for colinearity amongst the measures.

As large variations in prognosis rates have been reported among MCIs (Ward et al., 2013), we conducted a post-hoc analysis of variances in model parameters across and within groups. A Levene’s test with two factors – group (healthy, MCI, AD), and condition (Limbic SubNet, Whole Network) was used to assess homogeneity of variance of biophysical parameters.

#### 2.9.1 Estimation of model advantage over empirical FC and SC

To assess if there was any quantitative advantage in associating model parameters with cognition, compared to associating SC/FC with cognition, we performed the following: We conducted 100 bootstraps across subjects, for each iteration we performed two PLS analyses 1) biophysical parameters with cognition and 2) connectomes (SC and FC) with cognition. The 2 resulting distributions of obtained total covariances were saved.

The expected values for the covariances will differ between analyses because of scale differences. We corrected for this by conducting 100 permutations across subjects to build a null distribution for each analysis, each time summing covariances across all latent variables to take into account the whole spectrum of the covariance/relationship. The obtained total covariances were then corrected by the mean of its permuted null distribution.

The two resulting distributions of corrected covariances were compared by subtracting connectomes with cognition covariances from the biophysical parameters with cognition covariances. A difference distribution above zero would affirm that biophysical parameters correlate with cognition above and beyond SC and FC.

## 3. Results

### 3.1 Subject demographics, cognition, clinical grouping

Mean demographics, MMSE and IQ scores by group are shown in Table S4. Neuropsychological cognitive domain scores corresponded tightly with clinical diagnosis groupings; scores decreased consistently and significantly across the three clinical diagnosis groups (HC>MCI>AD; See Figure S5). Internal consistency scores of cognitive domain scores are shown in Table S5.

### 3.2 Model fitting and parameter estimation

Both the Limbic SubNet model and the Whole Network model generated good fits between the individual empirical and the simulated FC for each subject (Limbic SubNet: *r* = 0.57, *SD* = 0.075, 95% *CI* = [0.48, 0.65], Whole Network: *r* = 0.31, *SD* = 0.0453, 95% *CI* = [0.29, 0.33], mean *r*, *SD*, *CI* across subjects, for FCs generated with the optimal parameter values). The parameter search space showed a slow gradation towards a single optimal solution rather than multiple local maxima (See Figure S4). Thus, there is unlikely to be any appreciable interaction between any of the parameters. A systematic exploration of this has previously been shown using the same Deco et al. (2014) model (Schirner et al., 2018) and another similar model (Sanz-Leon et al., 2015).

Model fit did not differ significantly across groups, but was significantly higher for the Limbic SubNet than Whole Network model (2 factor ANOVA: group *F*(2,121) = 2.27, *p* = .15, model *F*(1,121) = 5817.77, *p* < 0.0001, group*model interaction: *F*(2,121) = 2.21, *p* = .11). We found that the 3 local parameters as well as global coupling correlated with model fit across subjects (See Table S6). However, conduction velocity did not influence model results in initial parameter explorations, thus time delays were not included in the model. Note that the effects we have reported are robust to using the SC weights normalized by track length. The optimal model parameters remained consistent when re-running the model with track length on two sample subjects.

### 3.3 Model correlates of cognition

We correlated model biophysical parameters with cognitive z-scores within the Limbic SubNet model, Whole Network model and the embedded Limbic SubNet (See Figure 2).

**Figure 2.**
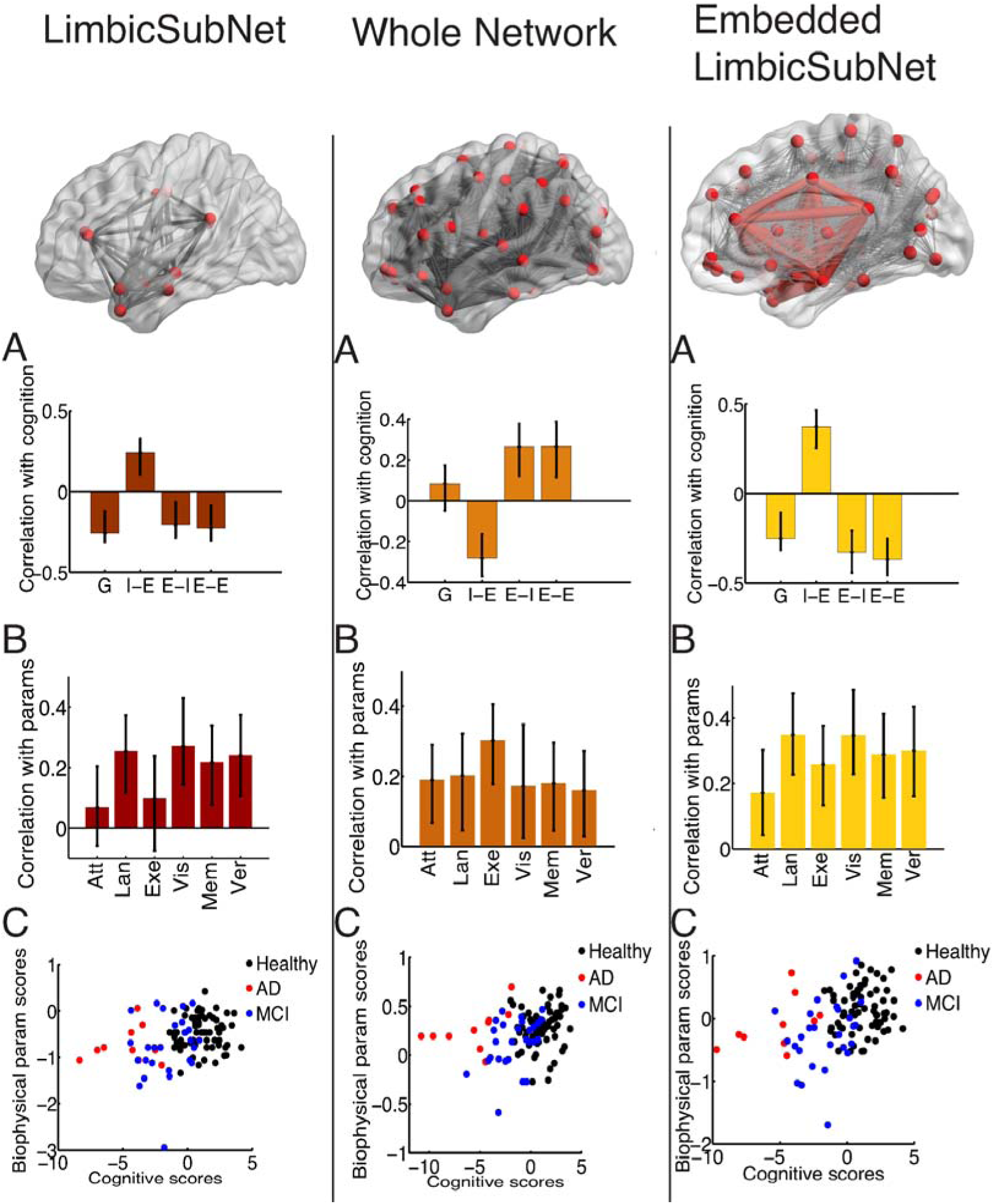
A) Correlation between biophysical model parameters and cognitive domain z-scores across all subjects for the Limbic SubNet, Whole Network model and the embedded Limbic SubNet. Biophysical model parameters are the optimal: global coupling, inhibition-excitation, excitation-inhibition, and excitation-excitation. Cognition domain z-scores include: attention (Att), language (Lan), executive function (Exe), visuospatial (Vis), memory (Mem), and verbal memory (Ver). B) The contributions of the cognitive domain scores to the cognition-parameter relationship. Confidence intervals on these correlations were obtained by bootstrap estimation. C) Individual subject cognitive domain scores and biophysical parameter scores, color coded across groups. These are the weighted sum of cognitive domain scores and biophysical model parameters per subject, respectively, and are similar to factor scores from factor analysis. The Embedded Limbic SubNet model was characterized as the model parameter discrepancy between the Limbic SubNet parameters and the Whole Network parameters.

We identified a single significant latent variable (LV) that characterizes the relationship between the biophysical parameters and cognitive domain z-scores for each of the models. Model parameters derived from the Limbic SubNet, the Whole Network, and the discrepancy between these two (the Embedded Limbic SubNet model) correlated with cognition on this LV (Limbic SubNet: *p* = 0.018, Whole Network: *p* = 0.01, Embedded Limbic SubNet: *p* < 0.001). The contributions of cognitive performance domains are shown in Figure 2B.

Within the Whole Network model, cognitive domain z-scores correlated negatively with inhibition, and positively with global coupling and excitation (excitation-inhibition and excitation-excitation). In contrast, within the Limbic SubNet, cognitive domain z-scores correlated negatively with global coupling and excitation (both excitation-inhibition, and excitation-excitation), and positively with inhibition.

We observed that the discrepancy between model parameters of the Limbic SubNet and the Whole Network correlates with cognitive performance (See Embedded Limbic SubNet model, Figure 2). Individuals with higher cognitive scores (i.e. healthy subjects) have a greater discrepancy in inhibition, and lower discrepancy in excitatory and global coupling parameters. Note that these results hold when correcting for subject demographics (age, non-English speaking background, sex, and education) via regression and using residuals for analysis.

### 3.4 Group differences in biophysical model parameters

Thus far we had compared biophysical model parameters with cognitive performance as a surrogate measure of disease severity. In order to examine whether parameters could also be used to differentiate between clinical diagnosis groups, we compared model parameters across clinical groups (healthy, aMCI, AD) and network (Limbic SubNet, Whole Network). Here, we did not observe group differences between parameters, nor an effect of network, but we did observe a significant group*network interaction (*p* = 0.004). See Table 1, and the same data in Figure 3, for optimal parameters and details by group.

**Table 1.**
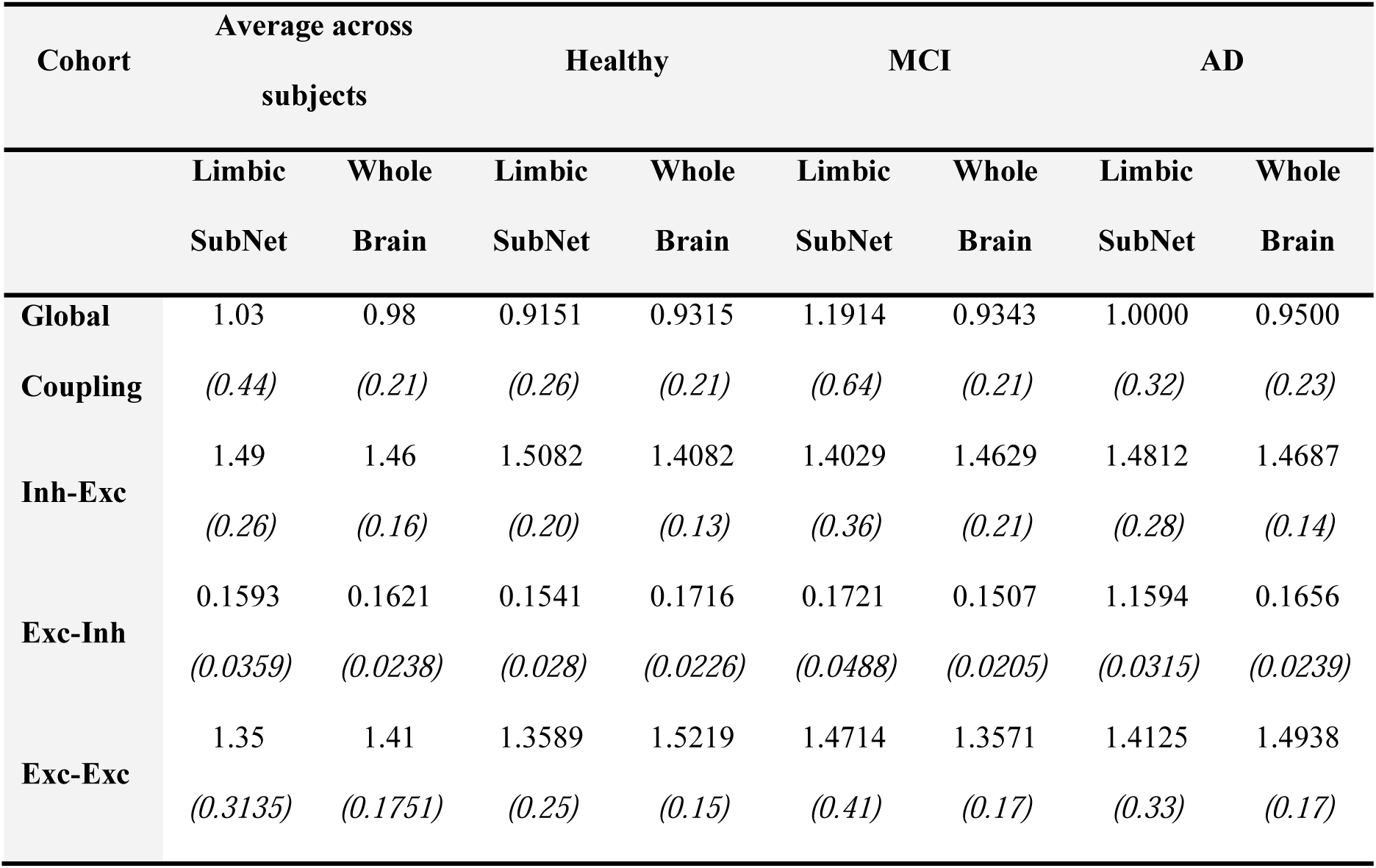
Mean (*SD*) of optimal biophysical model parameters per group and network

**Figure 3.**
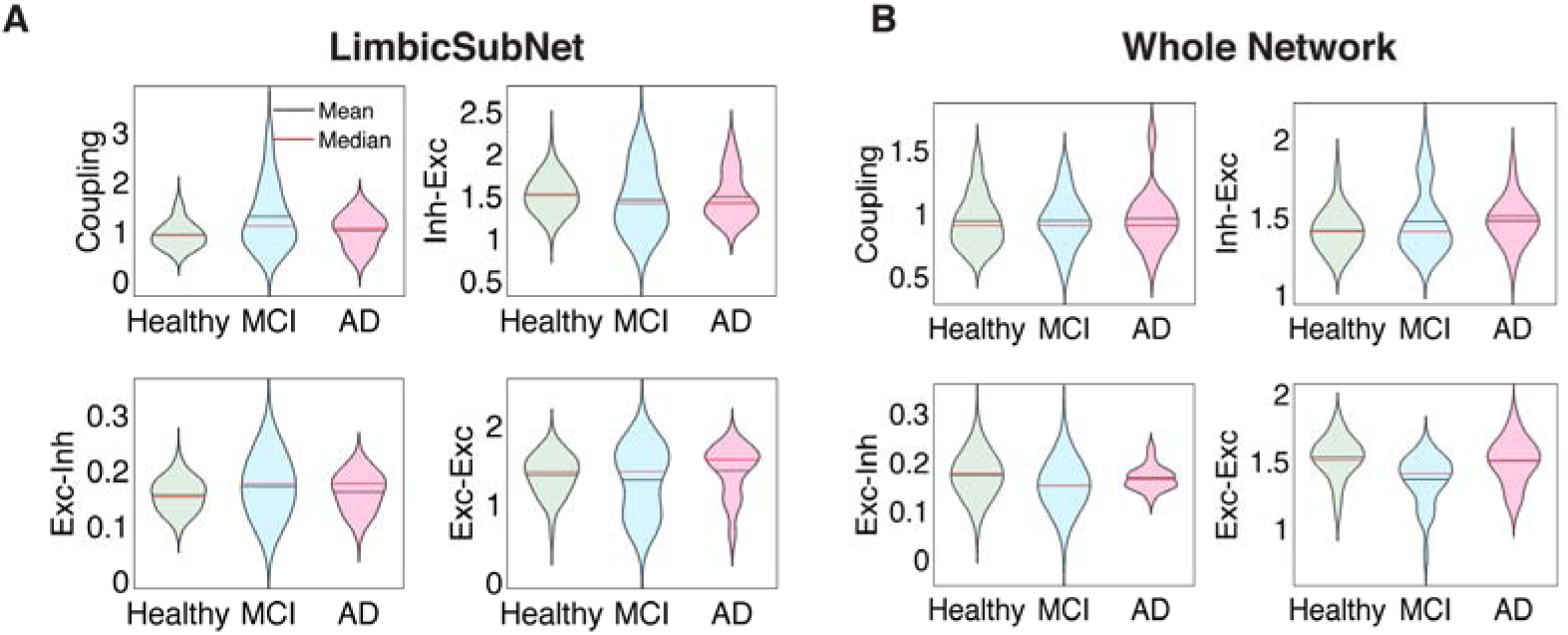
Variance of biophysical parameters across clinical groups for: A) the Limbic SubNet, where the greatest variance was observed among the MCI cohort, and B) the Whole Network, where the parameter variances were equal across groups.

Of note here (Table 1, Figure 3) was the large variance among the MCIs in the Limbic SubNet compared to healthy controls and ADs. A Levene’s test of homogeneity of variance for optimal values of each parameter was significant for group, network, and their interaction (except for the group*network interaction of inhibitory-excitatory parameter) (See Table 2).

**Table 2.**
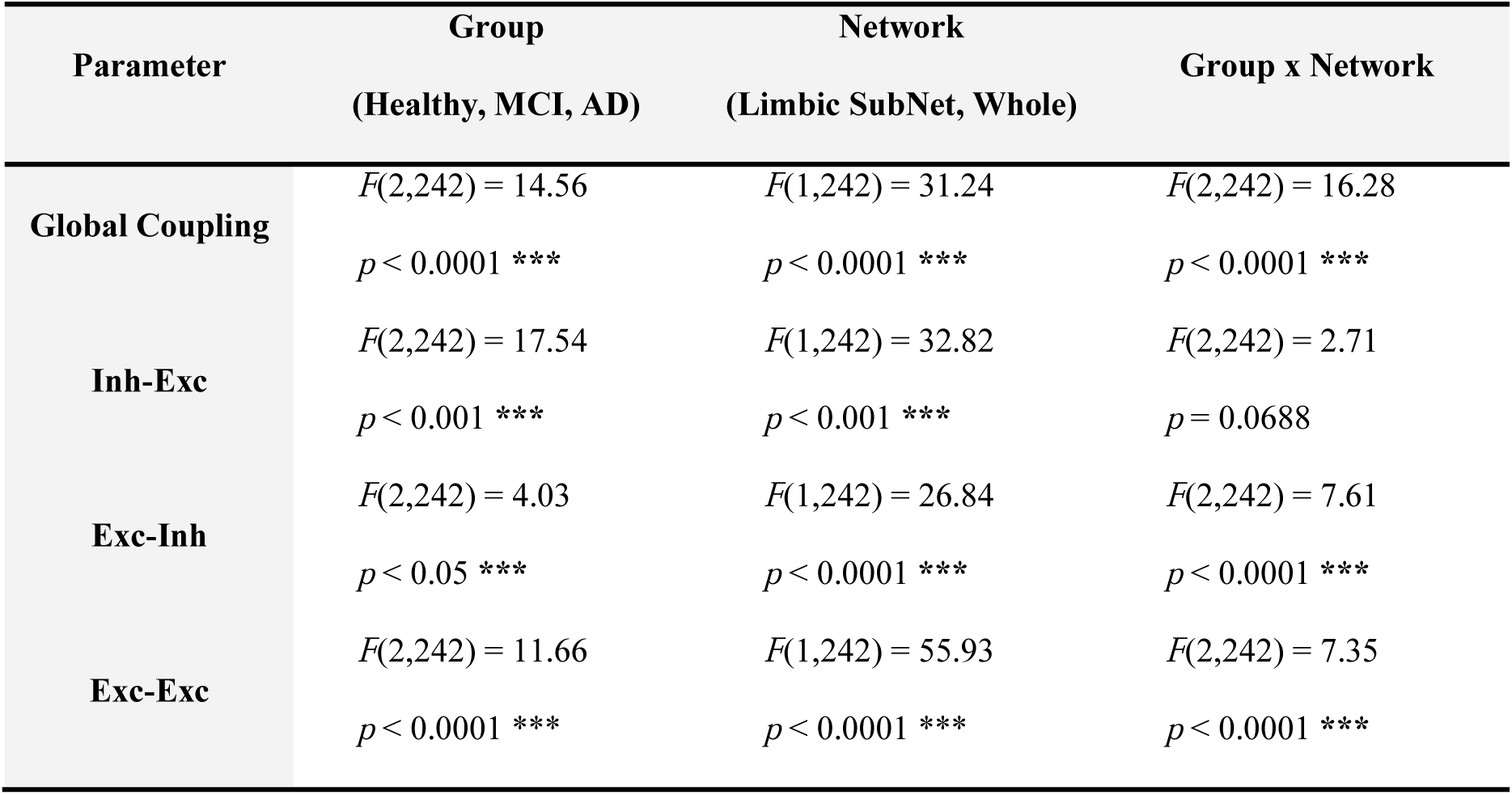
Levene’s test of variance in model parameters for group (healthy, MCI, AD) and network (Limbic SubNet, Whole Network), and their interaction

We hypothesized that the lack of differences between groups was due to the large variability in the aMCI group as well as the large differences in sample sizes between groups. In order to account for variable sample sizes that led to the unequal weighing of groups, we conducted a secondary analysis comparing the healthy and AD group, matching for age and sample size (N = 16 per group). Here, we reported a significant effect of clinical group (healthy, AD) (*p* = 0.003, *d* = 0.45) and network (Limbic SubNet, Whole Network) (*p* = 0.003, *d* = 0.73), as well as a group*network interaction (*p* = 0.02, *d* = 0.29).

### 3.5 Model advantage over empirical connectomes

Next, we compared our individualized patient models to empirical imaging data in their ability to predict clinical phenotype, for both the Limbic SubNet and Whole Network. We observed that: (1) the covariance between model parameters and cognition *exceeded* (2) the covariance between empirical connectomes and cognition (See Figure 4; Results shown here are for the Limbic SubNet model).

**Figure 4.**
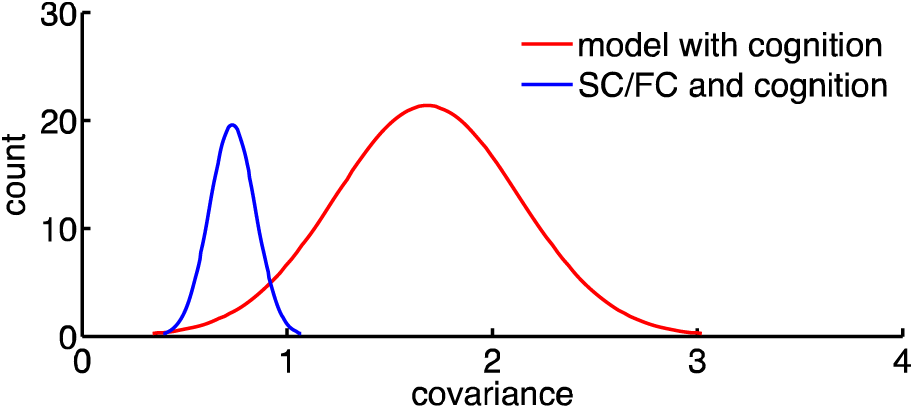
Covariance of biophysical model parameters with cognition outperforms covariance of connectomes (SC combined with FC) with cognition. Distributions shown are covariances from bootstrap resampling.

We bootstrapped covariances of (1) and (2) (See Methods), and subtracted the two to assert that model parameters with cognition significantly exceed connectomes (SC combined with FC) with cognition (See Methods). The mean of the difference distribution (the difference distribution was the connectomes with cognition covariance subtracted from model with cognition covariance) = 0.95, 95% *CIs:* [0.17, 1.70]. Thus, the difference distribution was reliably > 0.

See Supplementary materials for more details on reductions in empirical SC and FC weights (Figure S5), as well as graph measures across clinical groups. We also checked for SC-FC differences between groups, which were non-significant *F*(2,123) = 1.09, *p* = 0.338.

### 3.6 Subject motion

An ANOVA was conducted to compare framewise displacement between healthy controls, MCIs, and ADs to check whether subject motion was driving any differences between groups (Power et al., 2012). The result was non-significant (*F*(2,123) = 0.38, *p* = 0.68).

## 4. Discussion

We used a neurophysiological multi-scale brain network model (The Virtual Brain) to examine how individual optimal functioning points of local (inhibition/excitation) and global (long-range coupling) dynamics correlate with variability in cognition across healthy controls, amnesic Mild Cognitively Impaired (aMCI), and Alzheimer’s (AD) patients. The study is a compelling proof of concept that the modeling platform can be used to characterize an individual’s own network and local dynamics and has a clinical utility that can inform about the disease. We modeled, 1) the Limbic SubNet, which includes the regions of primary onset of the pathology (Van Hoesen and Solodkin, 1994), and 2) the Whole Network, which encompassed also the Limbic SubNet. We also characterized the embeddedness of the Limbic SubNet within the full network, which describes the influence of the larger network to the subnetwork.

We were interested in global long-range network changes (Delbeuck et al., 2003; He et al., 2008; Sanz-Arigita et al., 2010; Stam et al., 2007) as well as local excitatory/inhibitory dynamics on neural activity (Palop et al., 2007; Paula-Lima et al., 2013), as these changes have consistently been observed in AD. Quantitative metrics in our model reflect optimal levels of local excitation, inhibition, and global coupling for each individual. Such parameters have been linked to the emergence of important network features in health (Deco et al., 2009; Deco and Jirsa, 2012; Deco et al., 2014b), and performance in clinical conditions (Falcon et al., 2016b; Falcon et al., 2015). A link between model measures (e.g. excitation/inhibition) and function has been ascertained in computational models of AD in the past (de Haan et al., 2012; de Haan et al., 2017). Here, we showed these model parameters allow for the characterization of an individual’s own network and local dynamic changes in MCI and AD that reflect cognitive performance.

Specifically, we found that optimal levels of excitation, inhibition and global coupling in the individual subject’s model were associated with cognitive performance scores across six cognitive domains (attention, language, executive function, visuospatial, memory, and verbal memory). When disease severity was characterized by cognitive performance alone, there was a clear association between severity and biophysical model parameters (Figure 2). However, biophysical parameters did not differ when compared strictly across clinical diagnosis groups. This was likely due to the large variation in biophysical model parameters in the aMCI group (See Figure 3). The large variance in the optimal values of excitation/inhibition as well as global dynamics in the aMCI group was an important finding in our study, particularly in light of the heterogenic nature of this group in terms of phenotype and conversion. We conducted a secondary analysis in order to ascertain that biophysical parameters are nonetheless informative for clinical group. When matched for sample size and age, healthy controls’ biophysical parameters differed significantly from ADs’.

Particularly intriguing was the finding that the Limbic SubNet and Whole Network model showed opposing brain-behavioural patterns. We describe each below, and follow with an embeddedness explanation of this discrepancy.

### 4.1 Limbic SubNet

Within the Limbic SubNet, we observed that global inter-regional inputs and excitation correlated negatively with cognition, while inhibition correlated positively with cognition. The excitation/inhibition findings are consistent with the view that over-excitation is damaging to the system, leading to synapse debilitation, excitotoxicity, and subsequent cell death in AD (Palop et al., 2007; Paula-Lima et al., 2013), while inhibition serves to counteract its effects as a sort of compensatory mechanism to reduce vulnerability to excitotoxicity (Lapchak et al., 2000; Palop et al., 2007; Schwartz-Bloom et al., 2000; Velasco and Tapia, 2002; Zhang et al., 2007). On the other hand, the finding that global inter-regional coupling correlated negatively with cognition within the Limbic SubNet suggests that a limbic system where neuronal dynamics are driven by neuronal activity from other regions is maladaptive. This is consistent with shifts that have been observed in global dynamics in healthy aging (McIntosh et al., 2014), as well as maturation (Fair et al., 2009). Disease-related alteration in global coupling is in line with reports of connectome degeneration and of network reorganization that is governed by these long-range connections (Delbeuck et al., 2003; He et al., 2008; Sanz-Arigita et al., 2010; Stam et al., 2007; Supekar et al., 2008; Wang et al., 2007).

### 4.2 Whole Brain

The brain-behavioural relationships we observed in the Whole Brain model were quite different than in the Limbic SubNet. In the Whole Brain model, we found that excitation and inhibition were positively and negatively correlated with cognition, respectively. This suggests that the effects of excitation and inhibition are not straightforward and often unpredictable in a complex system like the brain. The positive cognition correlate of excitation that we found is in contrast with our findings in the Limbic SubNet (where cognition correlated negatively with excitation), as well as with the damaging effects of over-excitation that have consistently been reported (Busche and Konnerth, 2015; Celone et al., 2006; Dickerson et al., 2005; Gleichmann et al., 2011; Gleichmann and Mattson, 2010; Jones et al., 2016; Sperling et al., 2010). Interestingly, similar unpredictability of excitation on function in AD has been reported previously (de Haan et al., 2017). In a whole-brain computational model of AD, de Haan et al (2017) found that, contrary to their hypothesis, selective stimulation of excitatory neurons was actually beneficial for preserving network function. In the present study, excitation in the Whole Brain model was also positively linked to preserved cognition.

### 4.3 Embeddedness

Although the notion of embeddedness was first introduced within social-economic network sciences (Granovetter, 1985), it has more recently been used to characterize neural networks and to describe how the network exerts influence onto a subnetwork that lies within it (Misic et al., 2011; Vlachos et al., 2012). A subnetwork is embedded within a larger network if its behaviour or properties are affected by the outside system. Embeddedness can be characterized by the difference in dynamics between the network and the subnetwork. That is, the more different the dynamics, the greater the influence of the larger network on the subnetwork.

Here, we studied embeddedness in order to better understand the disparity in the cognition-parameter associations in the Limbic SubNet and the Whole Network model. We characterized the difference in optimal levels of functioning in the Limbic SubNet nodes and Whole Network, and therefore the dependency between the two (i.e. embeddedness), as a function of cognition. We showed that the dependency of these two sets of model parameters differ across severity, suggesting that the influence of the larger network on the Limbic SubNet varies in disease.

### 4.4 Variation in aMCI group

Of note was the striking variability in model parameters within the aMCI group. This is particularly interesting in light of the variability that exists in phenotype and conversion rates among aMCI patients. With an annual conversion rate of about 20% (Ward et al., 2013), only a certain portion of aMCIs will convert to AD within the next few years. We suggest that the high variability we found may reflect disparities in the classification of MCI and its subtypes. We propose that those aMCIs individuals that will be converters will have a biophysical parameter fingerprint closer to that of the AD group, while those that will not convert will have biophysical parameters closer to that of the healthy group. A follow-up study with MAS longitudinal multi-wave data will test the use of biophysical model parameters in the individual prediction of conversion.

### 4.5 Model parameters above empirical connectomes

Individual subjects’ model parameters were better predictors of cognition compared to empirical structural and functional connectivity. This is important, as our model adds value above and beyond SC and FC for making predictions about disease-related cognitive decline. The model identifies the key features of SC and FC that reflect the unique biophysical properties of the subject, thereby reducing the irrelevant noise inherent in the connectomes and capturing the most predictive representation of the empirical data. A related observation was made in a modeling study of stroke outcome (Falcon et al., 2016), whereby biophysical parameters at pre-therapy conditions were more strongly associated with long-term motor recovery than the patient’s physical features of stroke or their demographics.

### 4.6 Caveats and considerations

We note several possible limitations of our study. One concern is that the cognition-related differences in biophysical model parameters could be driven by non-biological factors. To reduce or correct for the influence of non-biological factors, a number of precautions were taken: 1) Quality control of SCs (See Methods), 2) analysis of model fits across disease severity, 3) analysis of SC-FC across disease severity. We conclude that neither model fit nor SC-FC relationships varied with severity and are thus unlikely to contribute to biophysical model differences in our study.

As the simulations were based on SCs, we conducted rigorous quality control of our matrices. Nonetheless, the validity of our SCs could be improved by using filtering methods that ensure the streamline weights more accurately resemble the underlying densities (Smith et al., 2013, 2015). In addition, the validity could also be enhanced by performing tracking upon the grey/white-matter interface (Smith et al., 2012). Whilst these methods were not employed, and we acknowledge the limitations of our widely-adopted normalization approach (Yeh et al., 2016), the current diffusion data do still benefit from being processed with the most advanced options currently available for the present acquisitions. These approaches have been shown to distinguish between normal ageing and neurodegenerative groups; Of note, a recent study which also employed fibre-orientation based estimations of SC, identified striking voxel-wise microstructural differences in MCI and AD patients (relative to controls) (Mito et al., 2018). Similarly, fMRI is a helpful tool for the identification of AD-related dysfunction at early stages of the disease (Sperling, 2011) in view of the synaptic changes that occur early in the course of the pathology, often before measurable clinical symptoms (Coleman et al., 2004; Selkoe, 2002), However, it is important to take into consideration that changes in rsfMRI in AD may be an effect of pathological neurovascular differences in the BOLD signal and hemodynamic response (Sperling, 2011). Studying AD-related changes using a neural model circumvents some of these issues.

We also chose a local neuronal population model that was sufficiently complex to capture the key features of interest (excitatory and inhibitory synaptic currents) but, as with all population models, embodying other abstractions. Future models could address more nuanced aspects of synaptic and circuit pathophysiology through the employment of more detailed multi-layer cortical models.

## Acknowledgements

The research was funded by the Weston Brain Institute (Weston Brain International Fellowship in Neuroscience) to J.Z., the NSERC grant (RGPIN-2017-06793) to A.R.M., and funding granted by the German Ministry of Education and Research (US-German Collaboration in Computational Neuroscience 01GQ1504A, Bernstein Focus State Dependencies of Learning 01GQ0971-5), the European Union Horizon2020 (ERC Consolidator Grant BrainModes 683049), Stiftung Charité/Private Exzellenzinitiative Johanna Quandt and Berlin Instititute of Health (BIH Johanna Quandt Professorship for Brain Simulation) to P.R., the Australian Research Council (CE140100007) to M.B and National Health and Medical Research Council (NHMRC) Australia Program Grant APP1093083 to W.W and P.S. The authors gratefully acknowledge the computing time granted by the John von Neumann Institute for Computing (NIC) and provided on the supercomputer JURECA at Jülich Supercomputing Centre (JSC) (www.fz-juelich.de, Grant NIC#8344 & NIC#10276 to P.R.).

